# The relationship of white matter tract orientation to vascular geometry in the human brain

**DOI:** 10.1101/2025.03.06.641646

**Authors:** Kurt G. Schilling, Allen Newton, Chantal M.W. Tax, Maxime Chamberland, Samuel W. Remedios, Yurui Gao, Muwei Li, Catie Chang, Francois Rheault, Farshid Sepherband, Adam Anderson, John C Gore, Bennett Landman

**Affiliations:** Vanderbilt University Institute of Imaging Science, Nashville, TN, USA; Department of Radiology & Radiological Sciences, Vanderbilt University Medical Center, Nashville, TN, USA; Department of Biomedical Engineering, Vanderbilt University, Nashville, TN, USA; Image Sciences Institute, University Medical Center Utrecht, Utrecht, The Netherlands; Cardiff University Brain Research Imaging Centre (CUBRIC), School of Physics and Astronomy, Cardiff University, Cardiff, UK; Department of Mathematics & Computer Science, Eindhoven University of Technology, Eindhoven, The Netherlands; Department of Computer Science, Johns Hopkins University, Baltimore, MD, USA; Department of Electrical Engineering & Computer Engineering, Vanderbilt University, Nashville, TN, USA; Sherbrooke Connectivity Imaging Lab (SCIL), Computer Science Department, Université de Sherbrooke, Québec, Canada; USC Stevens Neuroimaging and Informatics Institute, Keck School of Medicine of USC, University of Southern California, Los Angeles, CA, USA

## Abstract

The white matter of the human brain exhibits highly ordered anisotropic structures of both axonal nerve fibers and cerebral vasculature. Separately, the anisotropic nature of white matter axons and white matter vasculature have been shown to cause an orientation dependence on various MRI contrasts used to study the structure and function of the brain; however, little is known of the relationship between axonal and vascular orientations. Thus, the aim of this study is to compare the orientation between nerve fibers and vasculature within the white matter. To do this, we use diffusion MRI and susceptibility weighted imaging acquired in the same healthy young adult volunteers and analyze the alignment between white matter fibers and blood vessels in different brain regions, and along different pathways, to determine the degree of alignment between these structures. We first describe vascular orientation throughout the brain and note several regions with consistent orientations across individuals. Next, we find that vasculature does not necessarily align with the dominant direction of white matter in many regions, but, due to the presence of crossing fiber populations, does align with at least some white matter within each MRI voxel. Even though the spatial patterns of blood vessels run in parallel to several white matter tracts, they do not do so along the entire pathway, nor for all pathways, suggesting that vasculature does not supply/drain blood in a tract-specific manner.

Overall, these findings suggest that the vascular architecture within the white matter is closely related to, but not the same as, the organization of neural pathways. This study contributes to a better understanding of the microstructural arrangement of the brain and may have implications for interpreting neuroimaging data in health and disease.

## Introduction

The white matter of the human brain is composed of intricately organized structures, including nerve fibers and cerebral vasculature, which work in tandem to support the brain’s complex functions. Nerve fibers in white matter are composed of myelinated axons, which are bundled into tracts that connect different regions of the brain. These fibers exhibit a highly ordered, anisotropic arrangement [1, 2], and are typically classified as association pathways (connecting adjacent or distant regions within a hemisphere), projection pathways (connecting the cortex to the thalamus, brainstem, or spinal cord), or commissural pathways (connecting the two cerebral hemispheres) [1]. The location, connectivity, and orientation of many of these pathways have been well characterized in the literature [3], and are known to be crucial for facilitating communication across brain regions and supporting brain function, cognition, and emotion [4].

Alongside these neural pathways, the cerebral vasculature within the white matter is organized into a network of blood vessels whose structures are similarly anisotropic and facilitate delivery of oxygen to tissue. The vasculature of white matter is understudied relative to that of the cortex [5, 6], yet several studies have described the arteries and veins within the white matter in terms of location, origin, and orientation. For example, arteries have been categorized as subcortical (shortly ending in the cortex or immediately underlying white matter) or medullary (penetrating into the deep white matter) [7], with veins also classified similarly as subcortical (beginning in white matter and draining towards the pial surface) or medullary (draining deep into white matter) [8-10]. The vasculature orientation is known to traverse centripetally (towards the center) from the peripheral cortex towards the superolateral corner of the lateral ventricles. Because oxygen and nutrient delivery are necessary for proper function, the vasculature is often assumed to be ‘laid out with a definite relation to function’ [11] with several studies classifying vasculature in relation to known white matter pathways [5, 7, 10, 12]. However, the precise relationship between white matter and vasculature is understudied [12]. Specifically, it is unknown whether there is a relationship between the layout of white matter vessels and nerve fibers, and where they align and do not align throughout the white matter.

The highly anisotropic geometries of structures in the white matter cause orientation-dependent effects in a variety of MRI contrasts. For example, the diffusion of water molecules is hindered by the well-aligned axonal membranes and myelin sheaths [13], which is exploited by diffusion MRI and diffusion fiber tractography to map the fiber pathways of the brain [14]. Moreover, the orientation of white matter fibers with respect to the main magnetic B0 field affects the MRI signal for a range of MRI contrasts. Orientation effects of *white matter axons* have been reported for T2 and T2* relaxation [15, 16], T1 relaxation [17, 18], magnetization transfer [19], myelin water imaging [20], quantitative susceptibility mapping [21], and diffusion[22]. Similarly, orientation effects of *vasculature* have also been shown to affect both gradient echo and spin echo signals, influencing BOLD contrast [23], dynamic susceptibility contrast [24-26], and subsequently derived measures of blood flow and blood volume. However, many of these studies explicitly assume that blood vessels run in parallel with white matter fibers [23, 25, 26], an assumption that appears qualitatively reasonable but has not been validated.

Understanding the alignment between these two essential components is crucial for advancing our knowledge of brain microstructure and the interplay between neural pathways and their supporting vasculature. The potential alignment or misalignment between axons and blood vessels could have significant implications for interpreting neuroimaging data, particularly in the context of brain connectivity and neurovascular coupling. This relationship may offer further insights into how the brain’s vascular network is organized to meet the metabolic demands of different neural regions and pathways. This study aims to address this gap by comparing the orientation of white matter fibers and vasculature within the white matter using high resolution MR imaging techniques. We use diffusion MRI (dMRI) to map the orientation of white matter tracts and susceptibility-weighted imaging (SWI) to visualize the cerebral vasculature in the same healthy young adult volunteers. By analyzing the alignment between these structures across different brain regions and pathways, we seek to determine whether there is a consistent relationship between axonal and vascular orientations.

## Methods

Six healthy young adult volunteers (3 female, age range 24-32 years old) were scanned on both a 3T Philips Ingenia CX scanner equipped with high performance gradients (80 mT/m maximum amplitude; 200 T/ms/s slew rate) and a 32-channel head coil for diffusion data, and also scanned on a 7T Philips Achieva scanner equipped with a 32-channel receive coil for high resolution SWI data. This study protocol was approved by the Vanderbilt Institutional Review Board (IRB #020623). All research was performed in accordance with relevant guidelines/regulations, performed in accordance with the Declaration of Helsinki, and written informed consent was obtained from all participants in the study.

### Susceptibility weighted imaging

To acquire high resolution, high contrast images of vasculature, we used high field scanning (7T), multiple acquisitions, and performed super-resolution post-processing of susceptibility weighted images. We assume susceptibility variations reflect the distribution of deoxyhemoglobin in vessels [27, 28]. The susceptibility weighted contrast was acquired using a high-resolution, 2D gradient recalled echo (GRE) scan (FOV = 240 APx180 RL, CS-SENSE=6, flip angle = 55°, TE=16.2ms, TR=‘shortest’ 120 slices, total scan time = 7:58) reconstructed at 0.20×0.20 mm resolution with 1mm slice thickness. This sequence was repeated 4-6 times depending on scan time. Each set of magnitude and phase images was post-processed separately. First, phase images were high pass filtered using a Hanning filter, followed by applying a 4^th^ degree power function to the phase images, before multiplying by the magnitude image. The end result of these steps are anisotropic susceptibility weighted images.

Next, these images were super-resolved to 0.2mm isotropic image resolution using the SMORE algorithm [29]. Briefly, SMORE is an internally trained deep learning algorithm which exploits the different resolutions present in anisotropic volumetric images. It creates paired training data by first learning to simulate appropriately low-resolution patches (in our case, 0.20×1mm resolution) from the relatively high-resolution in-plane data (in our case, 0.20×0.20mm resolution). It then learns to approximate the mapping from low-resolution to high-resolution data from these paired samples, and finally applies the map to all cardinal through-plane slices. In this way, SMORE super-resolves the anisotropic volume without the need of external training data, solely using the relationship between in-plane and through-plane of the volume itself.

Finally, all repititions of the super-resolved isotropic SWI contrast were co-registered and averaged to increase SNR, accounting for the small amounts of motion between acquisitions. Additional imaging at 7T included a 3D T1-weighted dataset acquired using a high resolution 3D MPRAGE (TE = 2.6 ms, TR = 5.6 ms, flip angle = 7°, FOV = 256 × 256 × 180 mm, voxel size = 0.6 mm isotropic, CS-SENSE=8, TI = 1271ms, shot interval = 4500ms). These images were processed using the function *fast* from the FSL software library [30] to generate white matter, gray matter, and cerebrospinal fluid (CSF) masks.

### Vasculature segmentation and orientation

From the 0.2mm isotropic super-resolved SWI images, our goal was to both segment vasculature and to extract vascular orientation. These were performed following methods inspired by Sepehrband et al. [31], which were originally intended to quantify perivascular space. Briefly, images were first filtered using the adaptive non-local mean filtering technique [32] (implemented in Dipy software [33], with patch_radius=1 voxel, block_radius=5 voxels) to remove high frequency noise while preserving vascular signal intensities. Next, a Frangi vesselness filter [34] was applied to the images in order to detect tube-like structures in the SWI data. This filter is commonly used in medical imaging studies of vasculature, and works by analyzing the local eigenvalues of the Hessian matrix, which represents second-order image derivatives. The filter highlights regions with low derivatives in one direction (along vessels) and high derivatives in two orthogonal directions (perpendicular to vessels). The “vesselness” image was thresholded (using an empirically defined threshold) to segment both dark and light vessels that appear as cylinder-like structures. We note that the image was masked by the white matter mask obtained from the T1-weighted image to study white matter vasculature only. To further refine the mask, and minimize partial volume effects, both gray matter and CSF masks were dilated (3 voxels, 600um) and used to remove contrast that is at the boundary of these tissue types.

This filter additionally resulted in an orientation estimate for every voxel designated as a vessel. To facilitate comparisons with the lower resolution diffusion MRI data, the orientations were down sampled to 2mm isotropic by taking the average orientation of all vessels in each 2mm image patch. The final output from SWI processing was a 2mm isotropic image of the dominant vasculature orientation throughout the entire white matter.

For a population-level analysis, SWI orientations were propagated to MNI-152 space. This was performed by registering the SWI magnitude image to the ICBM 2009c Nonlinear Asymmetric T2w template [35] using ANTS antsRegistrationSyn tool, and applying the derived warp to the SWI orientations, rotated appropriately using the preservation of principal direction algorithm[36].

### Diffusion weighted imaging

Diffusion data were acquired using a pulsed-gradient spin-echo echo-planar-imaging sequence, and consisted of four *b*-values (*b* = 500, 1,000, 2,000, and 3,000 s/mm^2^), with 6, 15, 15, and 60 direction, respectively, and 11 *b* = 0 volumes (TE = 101 ms, TR = 6066 ms, slice thickness = 1.87 mm, flip angle = 78°, in-plane resolution = 1.8×1.8 mm). Data preprocessing included denoising (using both MPPCA [37] and Patch2Self [38]), and correction for susceptibility distortions, subject motion, and eddy current correction [39] using the PreQual preprocessing pipeline [40].

### Fiber orientation distribution

The diffusion-derived fiber orientation distribution (FOD) was derived using multi-shell multi-tissue spherical deconvolution [41] after multi-tissue response function estimation [42]. A rigid registration (using FSLs *epi_reg* command) was performed to estimate the transform from diffusion to SWI space, and FODs were transformed (using MRTrix3 *mrtransform* command) with appropriate FOD reorientation [43] and amplitude modulation [44]. Because this FOD represented a continuous distribution over a sphere, we wanted to extract just the local maxima of this function (i.e., peaks) in order to make direct comparisons with vascular orientation.

Peaks of the spherical harmonic function in each voxel were extracted using a Newton search algorithm along each of a set of pre-specified directions [45], with a maximum of three peaks per voxel. Peaks smaller than 10% the maximum value were discarded as false positives. This procedure resulted in an orientation estimate (or up to three orientation estimates) for every voxel in SWI space at 2mm isotropic resolution, representing the directionality of the white matter fiber populations.

### White Matter Regions and White Matter Pathways

Comparisons between vascular and fiber orientations were made in both white matter “regions” and white matter “pathways”. White matter regions were defined by the JHU ICBM White Matter Atlas [46, 47], which included non-overlapping labels for well-characterized white matter locations in the brain, typically in the deep white matter. To investigate vasculature orientation along entire pathways, fiber tractography was performed using the TractSeg [48]algorithm (in SWI space) from which 62 white matter pathways were segmented, including association, commissural, thalamic, striatal, and projection pathways. Multiple pathways may exist within the same MRI voxel, although potentially with different orientations [49]. Thus, these pathway-based analyses provide increased specificity over region-based analyses, where there are potentially many pathways and/or orientations in a region of interest.

### Orientation comparison

For both voxel- and region-based analysis, comparisons between SWI vascular orientation and diffusion fiber orientation were performed for every voxel at 2mm isotropic resolution. These are calculated using the inverse cosine of the dot product between the vectors, resulting in an angular deviation ranging from 0-90°. It is important to note that while vasculature had one dominant orientation, the white matter fibers within each voxel can contain up to three orientations. Thus, two comparisons were made for each voxel. First, the angle between vasculature and the *dominant fiber orientation* - i.e, the largest peak of the FOD, which represents the white matter direction with the largest apparent fiber density. Second, the angle between vasculature and the *nearest fiber orientation* - i.e., testing the angle between vasculature and all fiber orientations, and retaining the minimum angle. This tested not the dominant fiber orientation, but whether any orientation is relatively parallel to vasculature.

In contrast to the region-based analysis, the white matter pathway-based analysis contained only one dominant fiber orientation per voxel (i.e., the dominant direction of streamlines within a voxel). We performed along-tract analysis by computing the angular deviation between the single vascular orientation and the single fiber orientation along the length of the entire pathway, with each pathway segmented into 100 segments.

## Results

### Vascular orientation in individuals

**Figure 1** shows example SWI images for several subjects, in axial, sagittal, and coronal orientations, along with the derived vascular orientations. First, vasculature is clearly discernable as both dark and white tubular structures in all planes. Second, our image processing results in vascular orientation estimates that visually match those observed in the images. These quantitative orientations match qualitative observations in the literature. For example, the deep medullary vasculature is largely radially oriented, traversing to the lateral edge of the lateral ventricles, and traveling parallel to the gyral stalks within the sulcal folds. The vascular contrast is dominated by left/right oriented vessels in the level of the lateral ventricles and by superior/inferior oriented vessels throughout the superior corona radiata. Many vessels are seen to be oriented anterior/posterior throughout the frontal/pre-frontal white matter, and the orientations within the corpus callosum seem to be dominated by the vasculature/perivascular space along the ventricle, running along the length of the callosal fibers.

**Figure 1.**
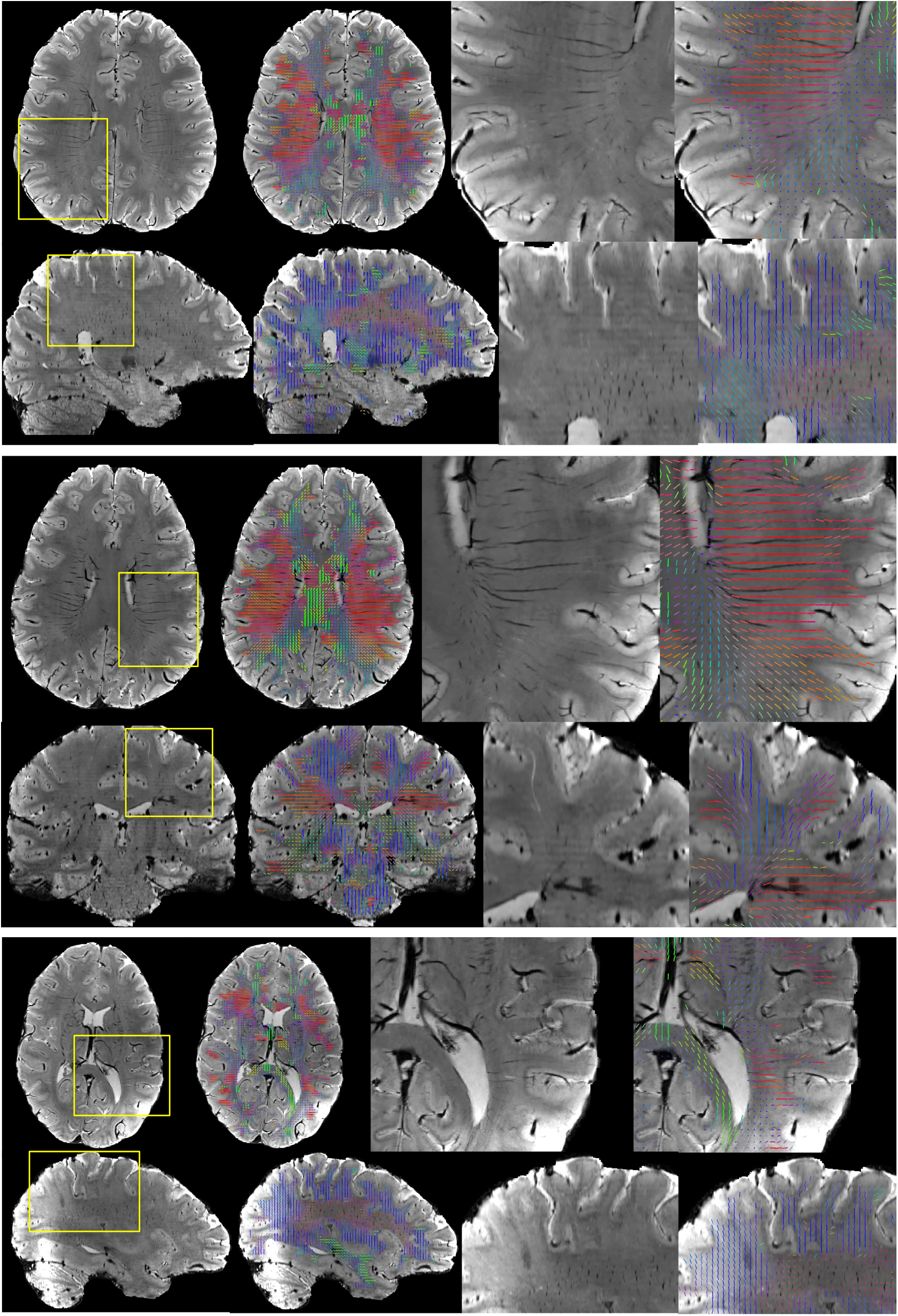
Vascular orientation in individual subjects. For three subjects, SWI images are shown in two orientations showing the full field of view and a zoomed-in region of interest. Overlaid vectors represent the dominant direction of vasculature in each 2mm region, and are color coded based on orientation, with red, green, and blue representing vasculature oriented in the left/right, anterior/posterior, and superior/inferior directions, respectively. Vasculature is clearly discernable in all imaging views, and orientations visually match the directionality observed in the images.

### Vascular orientation in the population

**Figure 2** shows the directionality of white matter vasculature averaged across our study population in MNI-152 space, along with the mean angular deviation across subjects as a measure of variability of vascular orientations. The population-level trends follow those observed at the individual level, with predominantly radial orientations from the lateral ventricle to the periphery of the brain, and vasculature aligning with the gyral stalks throughout. There is high homogeneity across subjects (low mean angular deviation) in the internal capsule and superior corona radiata as well as throughout gyri in all lobes of the brain. Orientation variability is higher in the corpus callosum, and lateral to the corona raditata in the area of the superior longitudinal fasciculi. There is a trend of high-low-high variability as we traverse anterior to posterior throughout the deep white matter.

**Figure 2.**
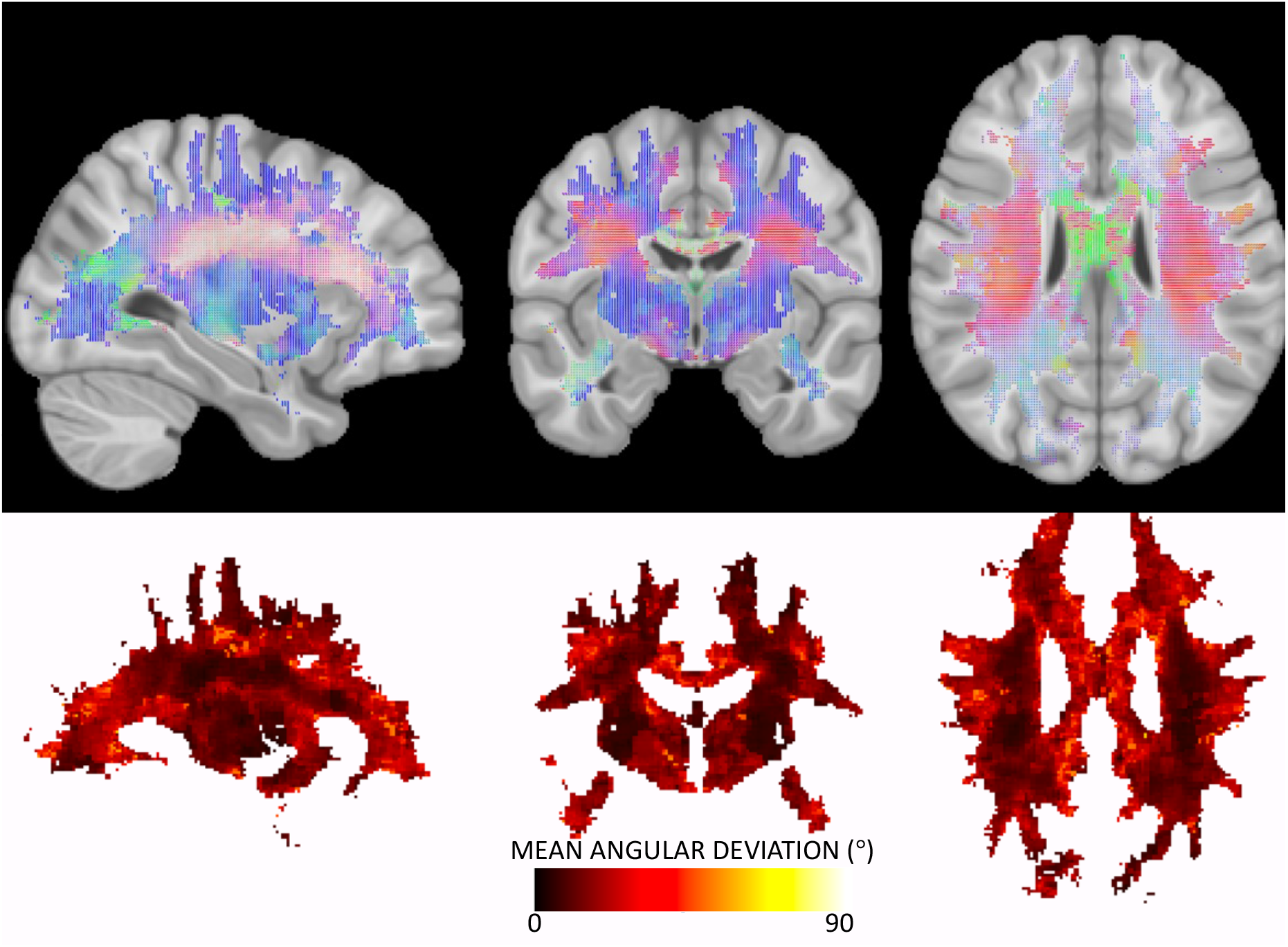
Vascular orientation averaged over the study population. Top row shows the population-averaged vector at each location in MNI152 space, with red, green, and blue representing vasculature oriented in the left/right, anterior/posterior, and superior/inferior directions, respectively. Bottom row show the mean angular deviation across all subjects, representing the average differences in orientation across subjects.

### Voxel-based alignment between vasculature and white matter orientation

**Figure 3** shows that vasculature typically lies nearly parallel with many white matter bundles but does not always align with the dominant white matter bundle in each voxel. For three example subjects, we show the structural image in three planes (top), the angular difference between vasculature and the *dominant* fiber orientation (middle), and the angular difference between vasculature and the *nearest* fiber orientation (bottom). There are many areas throughout the brain where vasculature is nearly perpendicular to large fiber bundles, but due to the presence and prevalence of crossing fibers, these vascular systems do lie nearly parallel to at least some fibers. This is quantified in **Figure 4**, where the angular difference is shown as histograms. Comparing vasculature to the most dominant axonal orientation in each voxel shows a bimodal distribution, with a mode of ∼18-26° of difference, but average difference of 41-48°, increasing for more orthogonal angles. When compared to the nearest fiber orientation, the angular differences are significantly reduced, with median values ∼28-34° with modes of ∼16-28°.

**Figure 3.**
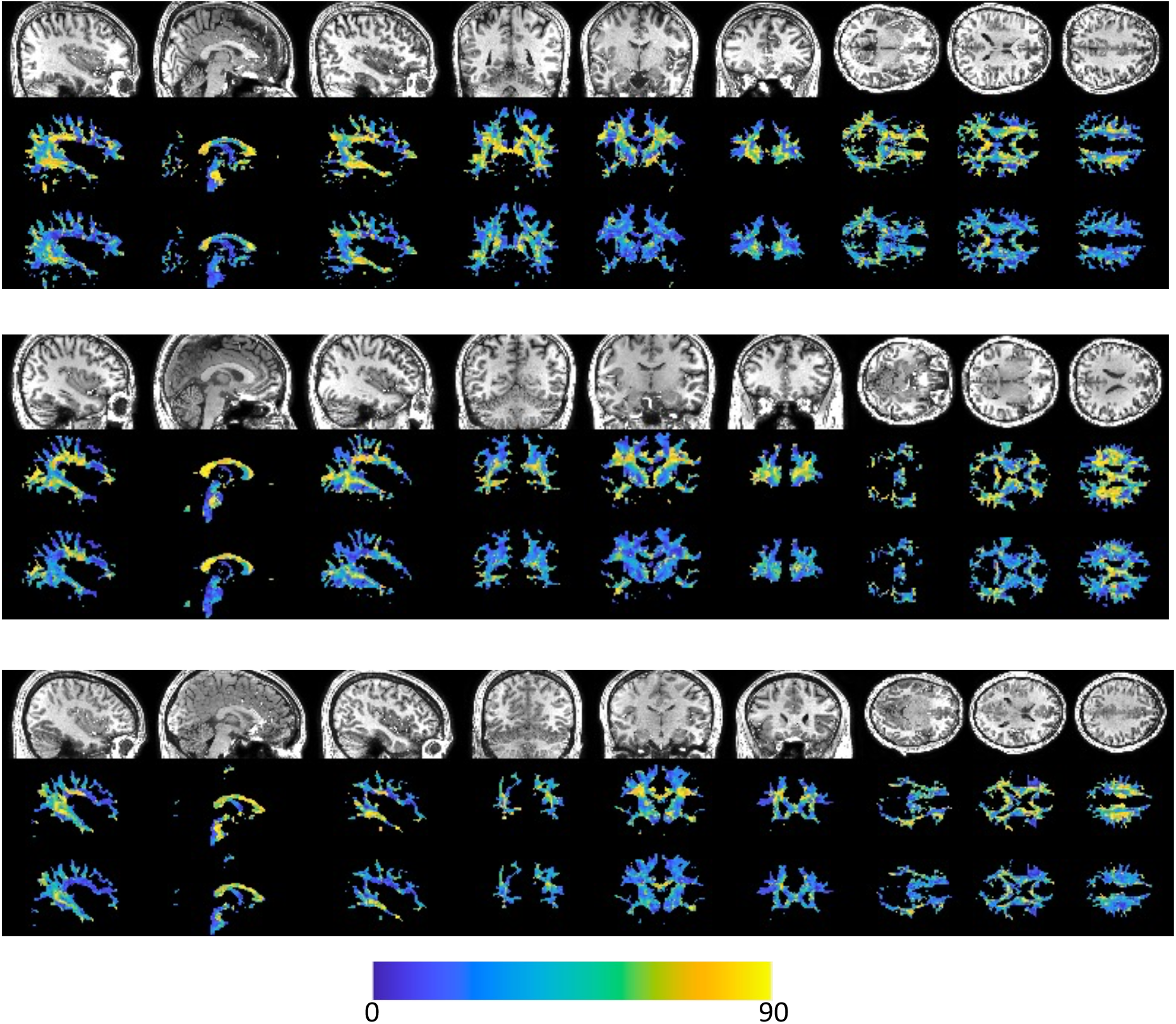
Alignment of vasculature and fiber orientation in the white matter. For three subjects, sagittal, coronal, and axial slices are shown for (top) the T1-weighted image, (middle) the angular deviation between vasculature and the dominant fiber orientation, and (bottom) the angular deviation between vasculature and the nearest fiber orientation. Vasculature does not always align with the dominant orientation, but typically parallels some white matter within each voxel.

**Figure 4.**
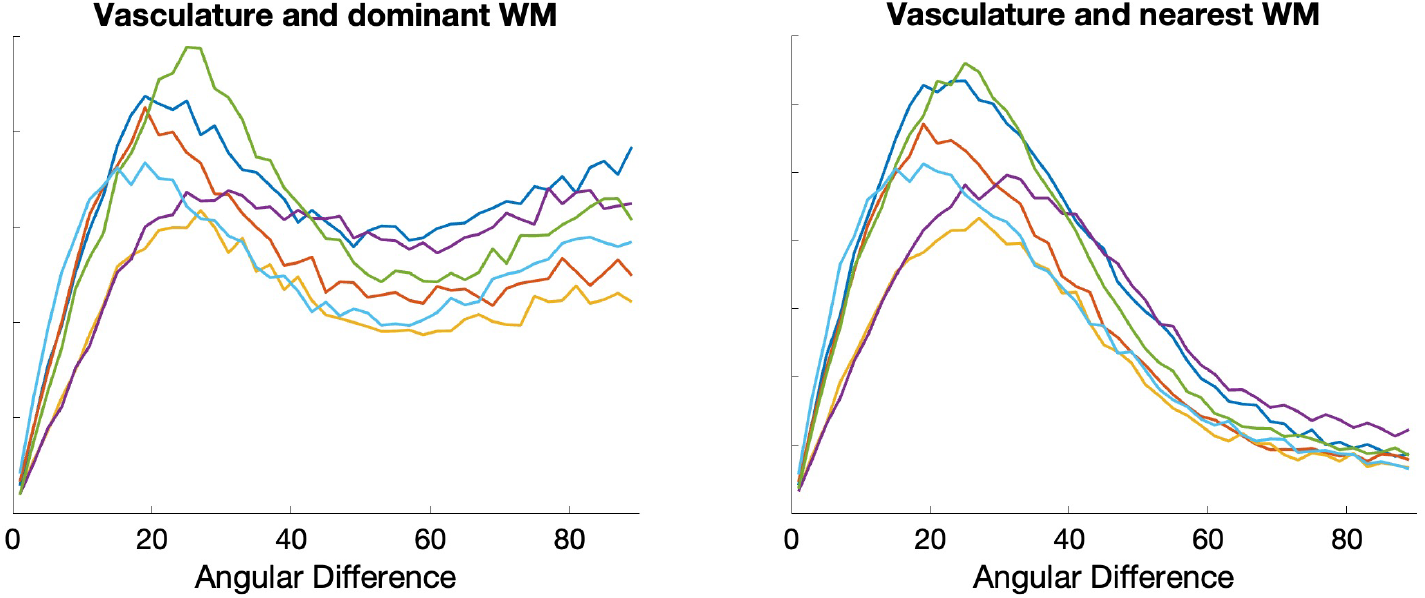
Histograms of angular differences between vasculature and fibers in the white matter. Plots show (left) the angular deviation between vasculature and the dominant fiber orientation, and (right) the angular deviation between vasculature and the nearest fiber orientation. Colors represent different subjects.

**Figure 5** shows similar results, but averaged in MNI space across the study population. Again, vasculature falls largely perpendicular or nearly parallel to white matter fiber structures, and throughout most of both deep and superficial white matter lies nearly parallel to at least some white matter pathways, with median and mean angular difference in MNI space of 28° and 30°, respectively.

**Figure 5.**
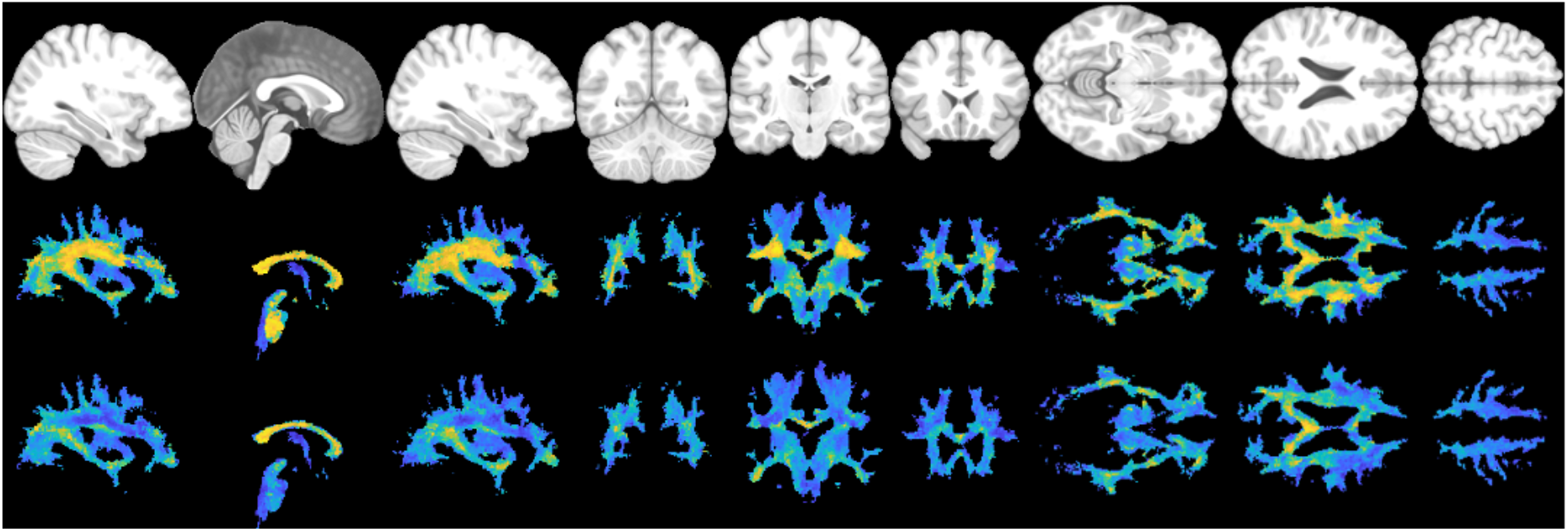
Alignment of vasculature and fiber orientation in the white matter, in MNI space. Sagittal, coronal, and axial slices are shown for (top) the T1-weighted image, (middle) the angular deviation between vasculature and the dominant fiber orientation, and (bottom) the angular deviation between vasculature and the nearest fiber orientation. Vasculature does not always align with the dominant orientation, but typically runs parallel to some white matter fibers within each voxel.

### Region-based alignment and misalignment

**Figure 6** shows boxplots for angular differences between vasculature and white matter fibers in deep white matter regions defined by the JHU white matter atlas. For each region, plots are shown of the angle between vasculature and both the dominant fiber orientation and nearest fiber orientation. The results mirror the quantitative observations in previous figures, with the largest differences between these two measurements observed in the corona radiata (anterior, superior, and posterior), and the superior longitudinal and superior fronto-occipital fasciculi. We note that these are averaged across relatively large regions with heterogenous differences in angles, and results tend toward the modes of the distributions shown in **Figure 4**.

**Figure 6.**
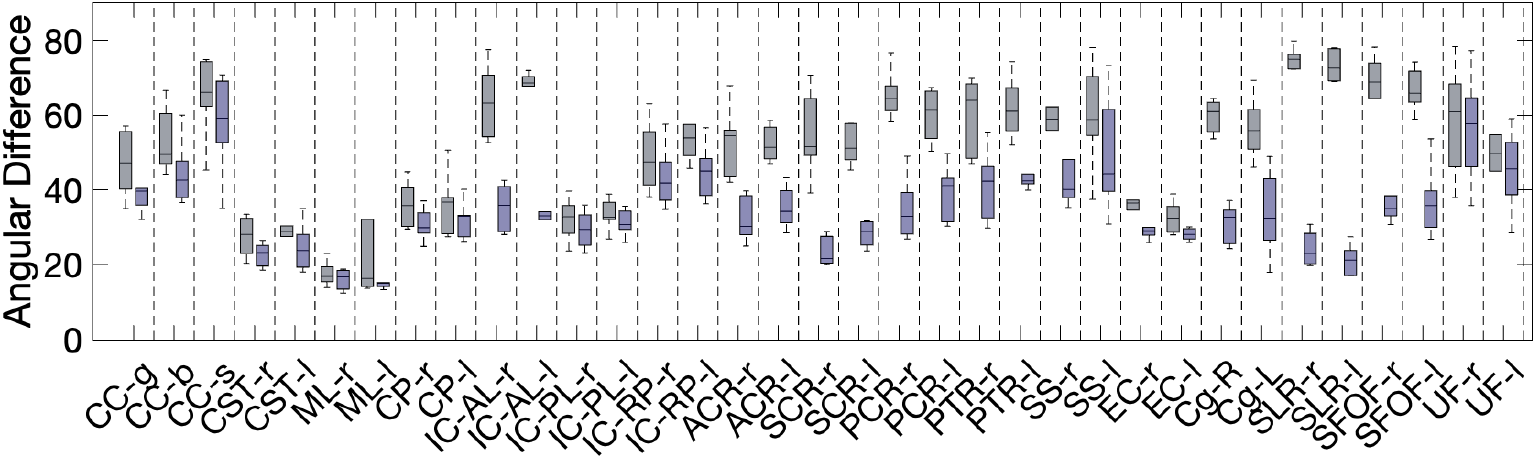
Angular difference between vasculature and fiber orientation in non-overlapping deep white matter regions defined by the JHU white matter atlas. Here, the average angular difference in each region is shown for (gray) the angular deviation between vasculature and the dominant fiber orientation, and (purple) the angular deviation between vasculature and the nearest fiber orientation.

### Pathway-based alignment and misalignment

Comparisons of vasculature and white matter orientation across entire pathways (which exhibit only one orientation per voxel) are shown in **Figure 7**. Boxplots (top) show the angular deviation averaged across each pathway, with pathways separated into association (green), commissural (red), thalamic (pink), striatal (magenta), and projection (blue). In general, vasculature lies most perpendicular to association pathways, and most tangential to striatal and projection pathways, with unique anterior-to-posterior trends in alignment seen across commissural, thalamic, and striatal pathways. Along-tract analysis (bottom) shows that vasculature-fiber alignment is not homogenous even within pathways. Several examples are shown, including a parallel-to-perpendicular-to-parallel alignment from the beginning to end of the arcuate fasciculi, or a parallel-to-perpendicular alignment along thalamic-premotor and optic radiations.

**Figure 7.**
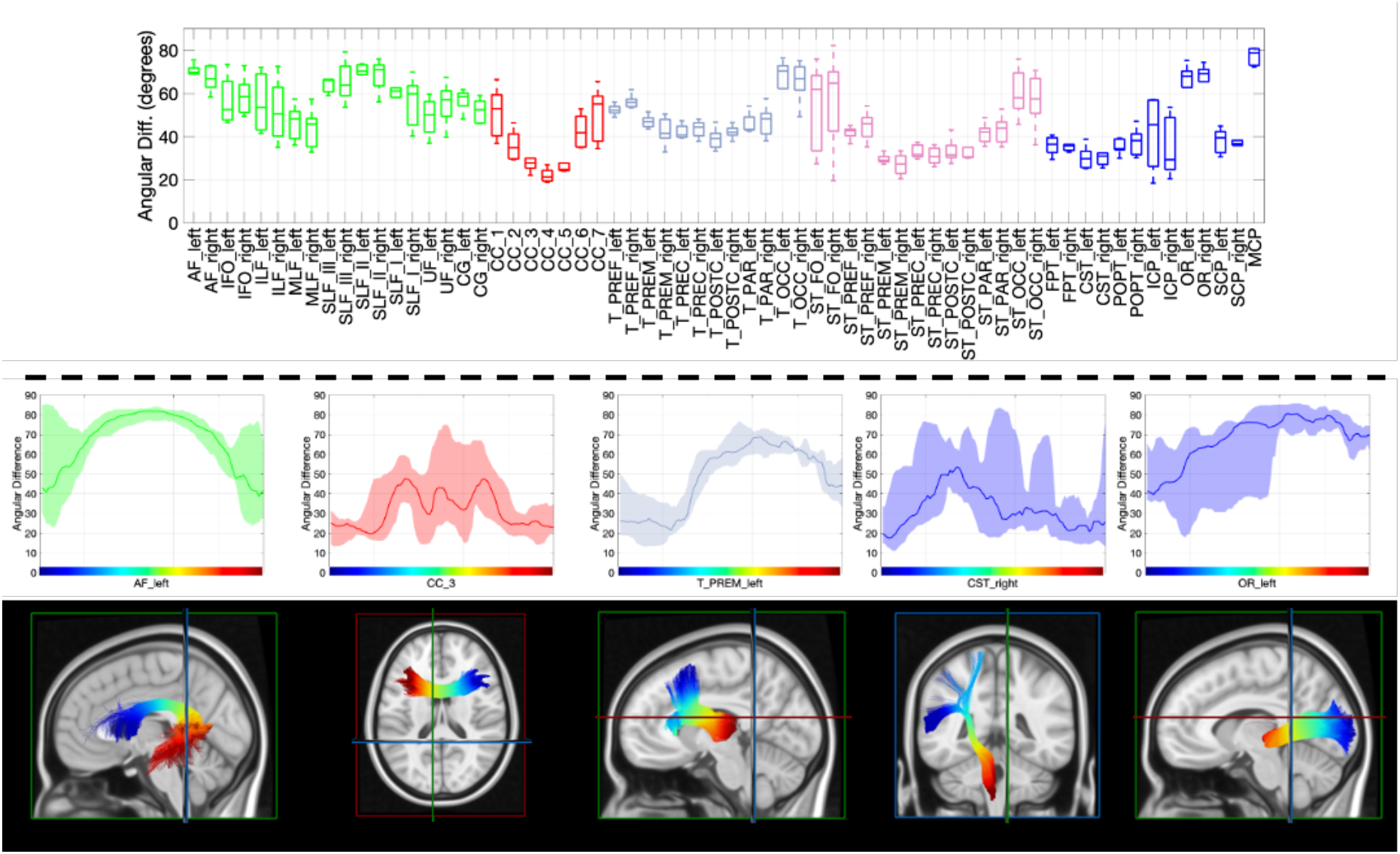
Angular difference between vasculature and white matter tracts (top) averaged along the length of the tract and (bottom) for every point along the tract from beginning to end. Note that plots are shown colored for association (green), commissural (red), thalamic (gray), striatal (purple), and projection (blue) fibers, and are plotted from beginning to end of pathway as designated in streamline-visualizations. Distribution shows the minimum and maximum across all subjects, highlighting the low variation across subjects along some pathways.

**Figure 8** shows streamlines overlaid on the SWI images where vasculature orientation is visible, for selected pathways. In all cases, arrows over the SWI images are oriented and colored to match the diffusion-derived streamlines. In the arcuate fasciculus, streamline terminations into the cortex at the frontal and parietal lobes are visible, running along gyri, and nearly perfectly aligning with vasculature. Alternatively, streamlines in the core of the pathway are entirely perpendicular to the vasculature. Thalamo-premotor pathway streamlines and vasculature are shown in the sagittal plane. Here, streamlines traverse laterally and dorsally from the thalamus (green-blue), with projections turning nearly and projecting superior (blue) to the premotor cortex. Vasculature in this plane is shown as obliquely oriented holes and finally superior-inferior oriented lines near the cortex, aligning with the pathway throughout its entire course. Finally, the optic radiations are shown in an axial plane. From the thalamus to the occipital terminations, the pathway goes from entirely perpendicular, to largely aligned with vasculature just posterior to the posterior end of the lateral ventricles, again highlighting heterogeneity in alignment within pathways.

**Figure 8.**
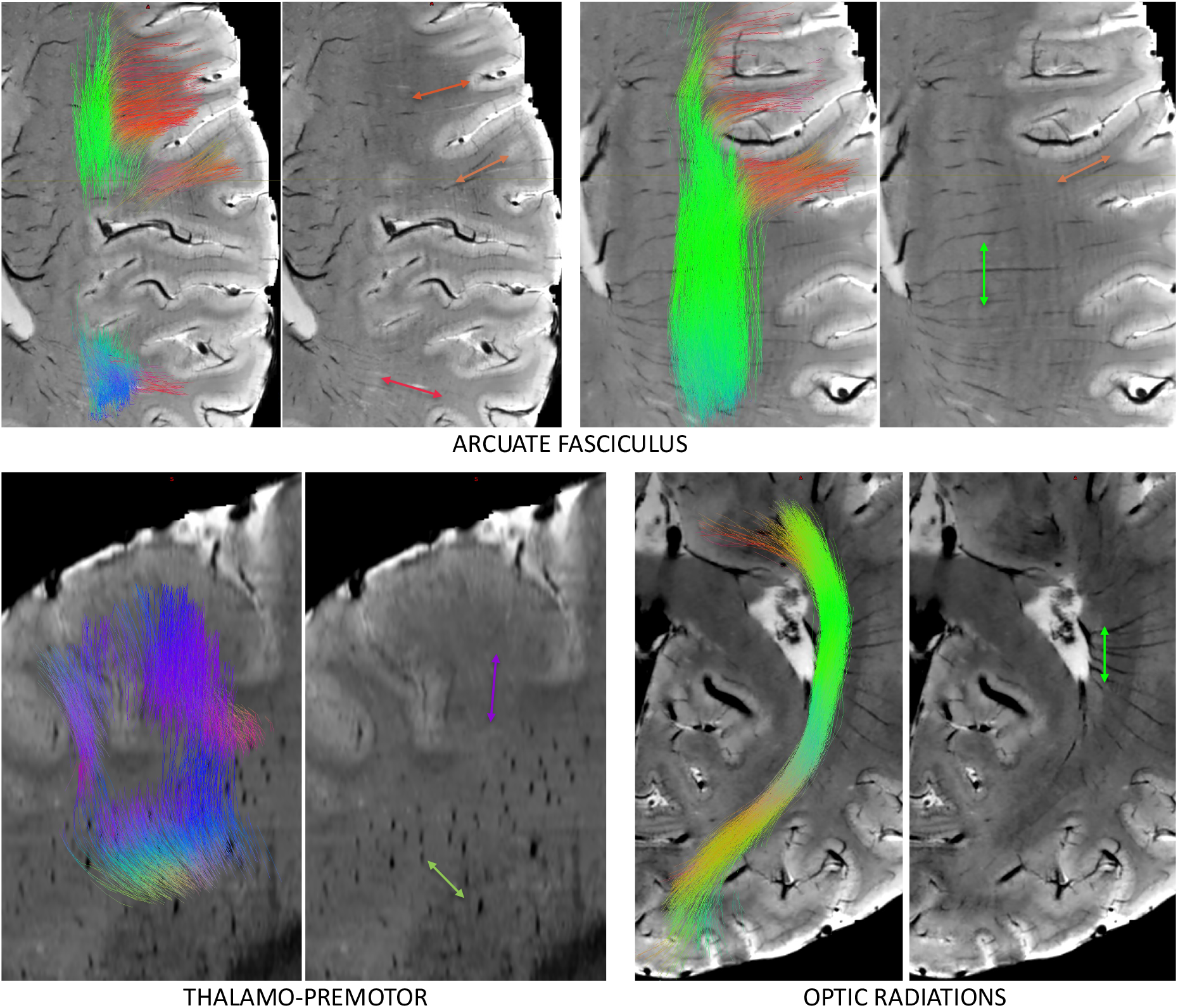
Alignment and misalignment between fibers and vasculature. For 3 selected pathways, including arcuate fasciculus (axial), thalamo-premotor pathway (sagittal) and optic radiation (axial), streamlines are shown overlaid on SWI. Streamlines are colored red, green, and blue, based on orientations in the left/right, anterior/posterior, superior/inferior directions. Arrows point in the direction of the streamlines and are colored with matching color at that location.

## Discussion

By acquiring high spatial resolution SWI images, and high angular resolution diffusion data, we compared vascular and nerve fiber orientations throughout the white matter on a voxel-by-voxel basis. In contrast to studies that assume a parallel arrangement between these two tissues, we find a more complex relationship where vasculature does not necessarily align with the dominant white matter direction in each voxel but does align with some bundles within each voxel. *Put another way, not all white matter bundles have corresponding vasculature along the direction of their axons*.

Based on existing literature and qualitative descriptions, we had hypothesized that most vasculature would align with white matter structures. In particular, the larger vessels of both veins and arteries in the white matter have been described as highly anisotropic [7, 9] and are visually consistent with many of the known white matter pathways of the brain [12]. However, this has not been assessed rigorously in a quantitative manner, which was the aim of this work. We found that many blood vessels (or sections of the vessels) do not align with any white matter axons, and those that do are not perfectly parallel, but rather often offset by ∼20°. This, though, does not necessarily disagree with previous literature. For example, two recent studies by Doucette et al. [25] and Hernandez-Torres et al. [24] investigating the effects of tissue orientation on dynamic susceptibility contrast show that empirically collected data matches simulations when one-half to one-third of vascular volume is comprised of vessels running parallel with fiber pathways. While we do not attempt to measure vascular volume, but rather perform analysis on a voxel-wise level, we find that a large fraction of voxels has aligning vasculature and nerve fibers.

### Vascularization of functional white matter pathways

As described in Smirnov et al., [5] most of the literature on brain vascularization describes cortical territories, with sparse quantitative descriptions of white matter regions as the “precise vascularization of most of the long association tracts is neglected”. This relationship between vasculature and the signaling pathways has direct clinical consequences where occlusions/infarctions of a given pathway may lead to functionally-specific deficits (e.g., posterior limb of the internal capsule and motor deficits, or inferior fronto-occipital fasciculus and semantic deficits). Here, we find that vessels do not appear to be pathway-specific. No pathway has parallel vessels along its entire length, and quite often traverse orthogonal to many white matter structures. This means that a vascular system does not supply/drain blood in a pathway-specific manner, and may be a form of redundancy - failure/occlusion of one specific segment will not lead to a complete failure of vascular function along an entire bundle.

However, the ability of white matter to perform signaling and housekeeping duties as a function of absent/impaired vasculature is unknown.

### Orientation and the MRI signal

The orientation of both white matter and vasculature has well-characterized effects and biases on a number of MRI contrasts, making it critical to understand the relationship between these tissues. It is well-established that both gradient-echo signals (and derived T2*) and spin-echo signals (and derived T2) depend on tissue orientation with respect to the B0 field [15, 16], with effects attributed predominantly to magnetic susceptibility of the myelin sheath surrounding axons. These effects have been convincingly demonstrated through simulations, and with empirical data, where the signal and derived contrasts have been directly related to the dominant direction of white matter in each voxel as determined by Diffusion Tensor Imaging (DTI). Digging deeper, recent studies using a tiltable coil and ultra-strong gradients have exploited diffusion and T2 contrast to find that these T2 effects are dominated by the extra-axonal signal (again, due to the myelin sheath) over the weaker, but still observable, effects due to the intra-axonal signal (also due to anisotropic microstructure, with magic-angle effects possibly due to oriented water molecules)) [15].

Beyond nerve fibers, other studies have indirectly investigated vascular anatomy by also observing a relationship in blood-related contrasts and the orientation, again, of *fiber* orientation derived by DTI - explicitly assuming a relationship between vascular and fiber orientations. For example, orientation effects of dynamic susceptibility contrast and blood-oxygenated-level-dependent (BOLD) contrast show orientation-dependent effects. In a recent study on BOLD fMRI, Viessman et al., [23] show a baseline signal orientation dependence consistent with either fiber and/or vascular orientation effects and also find BOLD fluctuation effects consistent with an effect due to blood oxygenation directly within the vessels. Different magnitudes of these effects were seen in different major white matter bundles, with pathways classified as baseline-dominated or fluctuation-dominated. While this heterogeneity across bundles is partially determined by the dominant orientation of the pathway relative to B0, it is likely also influenced by alignment/misalignment of vasculature with the bundle itself observed in our study.

Finally, vasculature in white matter has been shown to contribute to changes in diffusion tensor images and can affect interpretation and observed changes in derived parameters. Specifically, Sepherband et al., [50] show that perivascular space - the fluid-filled structure that accompanies vessels entering/leaving the cortex - has anisotropic, fast diffusing components that can have a substantial bias on voxel-averaged DTI measures. By incorporating a PVS into the model-fitting as a fluid-filled, anisotropic compartment, *aligned with the principal white matter direction*, these biases are reduced and give more specific insight into pathological or aging processes. Again, however, the PVS compartment was constrained to align with white matter nerve fibers based on qualitative observations of high-resolution anatomical scans. We show that this may not be the most appropriate constraint, although estimating directionality of a low contrast (high diffusing) component may be a challenge with conventional diffusion acquisitions. Regardless, mismatches between white matter fibers and white matter vasculature should be considered when fitting any quantitative model of tissue.

### Orientation within gyral folds

The pathways investigated in this study are major association, projection, and commissural pathways. One type of pathway that we did not investigate with tractography are short association fibers (SAFs) or superficial association fibers, the typically U-shaped systems immediately adjacent to the cortical surface, connecting nearby gyri in the same hemisphere. The reasons for not performing SAF analysis are due to challenges in the tractography process itself [51-53], and ambiguity/uncertainty in the definitions and location of these pathways from the lack of validation studies [54]. However, because of the direct parallel in nomenclature with vasculature (superficial draining veins or superficial medullary arteries), it is worth discussing the relationship between these two structures. The superficial vasculature runs perpendicular to the cortical surface, through the sulcal walls, curving sharply (sometimes at right angles) to generally follow the direction of white matter within gyral folds, typically parallel to the gyral stalk [1]. This is nearly identical to descriptions of white matter axons entering the cortex, running tangential to the cortical boundary and turning sharply once reaching the cortex [55]. Our results suggest that the vasculature and white matter share similar orientations in the superficial white matter areas (low angular deviation in Figure 1, alignment with streamlines in Figure 8). Future studies will attempt to trace individual cylindrical contrast within SWI images (veins/arteries) and make direct streamline-to-streamline comparisons with traditional tractography, to query whether these systems stay in alignment, or query where and when these systems first diverge. It is possible there is no relationship other than the radial organization of vasculature to minimize distance from the periphery (cortex) to the deep white matter.

### Limitations

There are several limitations to this study that must be acknowledged. First, is the limited sample size. Despite this, results generalized across the scanned cohort, and primary analysis is based on intra-individual comparisons. Second is limited vasculature sensitivity due to image resolution. Our images are acquired at a very high resolution (0.2×0.2×1mm) using state-of-the-art imaging and super-resolved to 0.2mm isotropic resolution using state-of-the art image processing, resulting in some of the highest resolution whole-brain in vivo images in the literature today. However, sensitivity to small vessels is still limited, but we note that vessels smaller than the voxel sizes are still detectable due to susceptibility differences of the blood and due to the image processing (Frangi filter) sensitivity to small changes in contrast. Further, the anisotropic components of vasculature that we are interested in are dominated by the larger vessels. Regardless, our conclusion of “not all white matter bundles have corresponding vasculature along the dominant direction of their axons” is valid only for vasculature that our image contrast is sensitive to.

Because of challenges at 7T, there are B0 artifacts at the limits of the field of view, most often in the temporal lobe or brainstem areas. For this reason, most descriptions are focused on cerebral vasculature (not including brainstem), although several of the pathways-of-interest travel through both brainstem and temporal lobes. These artifacts lead to changes in contrast that will be detected by the image processing filters, resulting in false positive detection of vasculature in these regions. For this reason, or empirically determined threshold favored specificity over sensitivity when detecting vessels. Again, despite this, the derived orientations well-matched those observed directly in the images. In contrast, it is possible that even higher field strengths, with 9.4, 11.7 [56], and future 14T [57] in vivo machines may enable more detailed mapping of white matter fibers and white matter vasculature.

## Conclusion

The white matter of the human brain exhibits highly ordered anisotropic structures of both axonal nerve fibers and cerebral vasculature. Understanding the relationship between these tissues can provide insight into developmental and clinical trajectories, and can help inform interpretation of MRI signal. While the anisotropic nature of each has been studied individually, little is known about the relationship between their orientations. This study compares the orientation of nerve fibers and vasculature within the white matter using diffusion MRI and susceptibility-weighted imaging in healthy volunteers. We found that while vasculature sometimes aligns with white matter tracts, this alignment is inconsistent across brain regions and pathways. These findings suggest that although the vascular architecture is related to neural pathways, it is not organized in a strictly tract-specific manner.

## Acknowledgements

R01EB017230 (Landman), K01EB032898 (Schilling), R01MH123201 (Gore, Landman), R01NS078680 (Gore), R01NS113832 (Gore). Wellcome Trust 215944/Z/19/Z (Tax). For the purpose of open access, the author has applied a CC BY public copyright licence to any Author Accepted Manuscript version arising from this submission. Dutch Research Council (NWO).

Grant Number: OCENW.M.22.352 (Chamberland).

## Data Availability

The datasets used and/or analysed during the current study available from the corresponding author on reasonable request.

